# A critical role for spike synchrony in determining steep 1/f slopes in the setting of bursting EEG patterns

**DOI:** 10.1101/2022.05.12.491724

**Authors:** Justin D. Yi, Yama Akbari

## Abstract

EEG signals are commonly analyzed in the frequency-domain. Analysis of the power spectrum commonly demonstrates a reproducible fall-off of high frequencies, sometimes referred to as the “1/f slope”. The 1/f slope comes about from a linear fit of the power spectral density in log-log form and takes typical values between (minus) 0–4. This slope has been recently hypothesized to correlate with the ratio of excitatory to inhibitory synaptic weights. However, we demonstrate models of excitatory-inhibitory (E/I) balance only explain small values of 1/f slope (< 2). We seek to construct a model to explain large slopes (> 2), which have been reported under conditions of anesthesia. Using simulations of clustered spike pulses, we find that pulse widths on the order of 1-10 ms result in a transition to steep 1/f slopes between 3–4. This trend also holds when measuring the 1/f slope from a spiking recurrent network model in a synchronous regime. We conclude that steep 1/f slopes result from synchronous spiking in bulk microcircuit activity. This suggests interpretations of steep 1/f slopes should consider spike synchrony in addition to other factors like E/I balance.

## Introduction

Analyzing electroencephalogram (EEG) data in the frequency-domain is ubiquitous. The power spectrum is a natural tool for EEG since there exists canonical frequency-bands where EEG activity is most active, providing a snapshot of EEG activity. However, purely analyzing EEG in the frequency-domain has its pitfalls which have recently been questioned. Without proper consideration for changes in aperiodic signal contributions which may permeate the power spectrum at all frequencies, measuring changes in specific frequency-band power can lead to the false positives [Donoghue et al., 2020a, 2021]. Even time-frequency methods like wavelet transforms, which seem at first like a compromise between time- and frequency-domain methods, run the risk of classifying spurious phase-amplitude-coupling between frequency-bands, when really such phenomena are better described by non-sinusoidal waveform shapes [Cole et al., 2017, Cole and Voytek, 2017, 2019]. In short, EEG analysis without explicit consideration of time-domain phenomena which generate its features can be misguided.

But, computationally understanding time-domain patterns in a meaningful and reproducible way requires defining (potentially automated) summary features [Fulcher and Jones, 2017]. Yet the challenge does not end with identifying and measuring features, which can already be a computationally expensive task. It is well established that even with a large set of well-defined summary features from neural time-series, the space of possible mechanistic models is vast [Marder and Taylor, 2011, Gonçalves et al., 2020, René et al., 2020]. Based on these limitations, what is needed is a metric that has roots in mechanism, summarizes the time-domain dynamics, but is as easy to calculate as other commonly used frequency-domain metrics like sub-band power, etc.

One candidate is the so-called “1/f slope.” 1/f slope is the name given to the characteristic linear fall-off of high frequencies in the EEG power spectrum when plotted in log-log scale. The power spectrum can be split into periodic and aperiodic activities [Donoghue et al., 2020b], where the 1/f slope is thought to comprise the aperiodic component. Gao et al. [2017] pointed out that the 1/f slope varies systematically with the ratio of excitatory to inhibitory synaptic currents from neurons firing asynchronously. This interpretation of the 1/f slope has been used to infer the balance of excitatory (E) and inhibitory (I) currents in the cortex for a variety of states of health, age, and disease [Robertson et al., 2019, Trakoshis et al., 2020, Schaworonkow and Voytek, 2021]. The 1/f slope is simple to calculate and has potential to illuminate details of the neural circuits that usually require invasive methods to probe.

Despite the recent promise of the excitatory-inhibitory (E/I) hypothesis, there has been an unaddressed mismatch between theory and observation surrounding the range of possible 1/f values. For instance, it has been argued that the 1/f slope becomes steeper (i.e. has larger absolute values) for smaller E/I ratios assuming Poisson spiking statistics [Gao et al., 2017]. The local field potential (LFP) has been approximated as the sum of E and I currents. From this LFP, the power spectral densities (PSD) can be calculated to find the 1/f slope. Since the 1/f slope depends on the E/I ratio, it thus attains a maximal value equal to the slope of the I PSD alone, i.e. when *I* ≫ *E*. Assuming the I synaptic conductance triggered by a single pre-synaptic inhibitory spike can be expressed as the difference of exponentals,

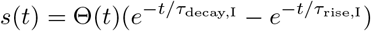

where *τ*_decay_ ≫ *τ*_rise_ and Θ(*t*) is the Heaviside step function, the resulting power spectral density, *S*(*ω*), can be approximated by a Lorentizian

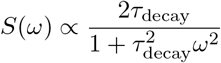

[Garcia et al., 2016, Theodorsen et al., 2017]. Thus, the maximal 1/f slope achievable by this linear model is (−)2 since

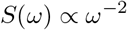

as *ω* → ∞. For a simulation of a recurrent network, Trakoshis et al. [2020] found that (for slopes measured at 30 – 100 Hz) the 1/f slope asymptotes around 2.0 to 2.5. Note, only for large frequencies (80 – 500 Hz) and where the condition *τ*_decay_ ≫ *τ*_rise_ is not satisfied, the power spectrum does fall-off like

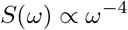

as detailed in Miller et al. [2009]. However, experimental observations show that 1/f slopes can reach extreme values like 3.0 or 4.0 in the 30 – 60 Hz range [Gao et al., 2017, Schaworonkow and Voytek, 2021], which are not explained by simulations in Trakoshis et al. [2020] nor by the linear superposition of E and I currents presented in Gao et al. [2017]. Barring the argument by Miller et al. [2009], what can account for these extremely steep slopes measured at low frequencies?

A step forward may be found in understanding the conditions which lead to steep slopes, namely the requirement of anesthesia. Gao et al. [2017] found propofol induced extremely steep 1/f slopes, and Colombo et al. [2019] correlated steeper 1/f slopes with increasing degrees of unresponsiveness/unconsciousness from anesthesia. General anesthesia has been shown to cause cortex-wide synchronized bursting selectively in the layer V pyramidal cells [Bharioke et al., 2022] resulting in a state clinically referred to as burst-suppression [Steriade et al., 1994, Shanker et al., 2021]. While bursting is grossly synchronized, there has been local variability in burst-onset time noted [Lewis et al., 2013]. In rodents, cortex-wide propagation of activity from a focus was observed with mean spreading times of about 54 ± 25 ms [Ming et al., 2021]. Therefore, the key to understanding the generation of steep 1/f slopes may lay in studying the effect of including highly synchronized spike timings, rather than assuming Poisson spike arrival statistics.

## Results

### 1/f slope as a function of E/I balance and spike timing

One possibility to explain large 1/f slopes is strong deviation of the stimulating spike trains from Poisson statistics. To test this, we design a spiking stimulus which appears to be Poisson only at time scales much smaller or much larger than the subthreshold membrane time constant of our model neuron, *τ_m_* = 15 ms. In the middle time scales, it looks like clustered pulses of spikes which cannot be approximated by a homogeneous Poisson arrival process.

Consider the transmembrane current resulting from *N* spikes arriving each independently at times {*t*_1_, *t*_2_, …, *t_i_*…, *t_N_*} within some interval *σ*_0_ ≪ *τ_m_* defined by a Gaussian centered on times *t_j_* sampled from a latent Poisson process with rate λ (see Fig. 1A). This will be called a Gaussian Packet Model (GPM). The spike trains of the GPM in the limit of *σ*_0_ ≪ *τ_m_* can be approximated by

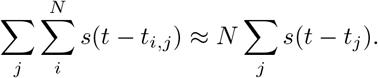

**Figure 1:**
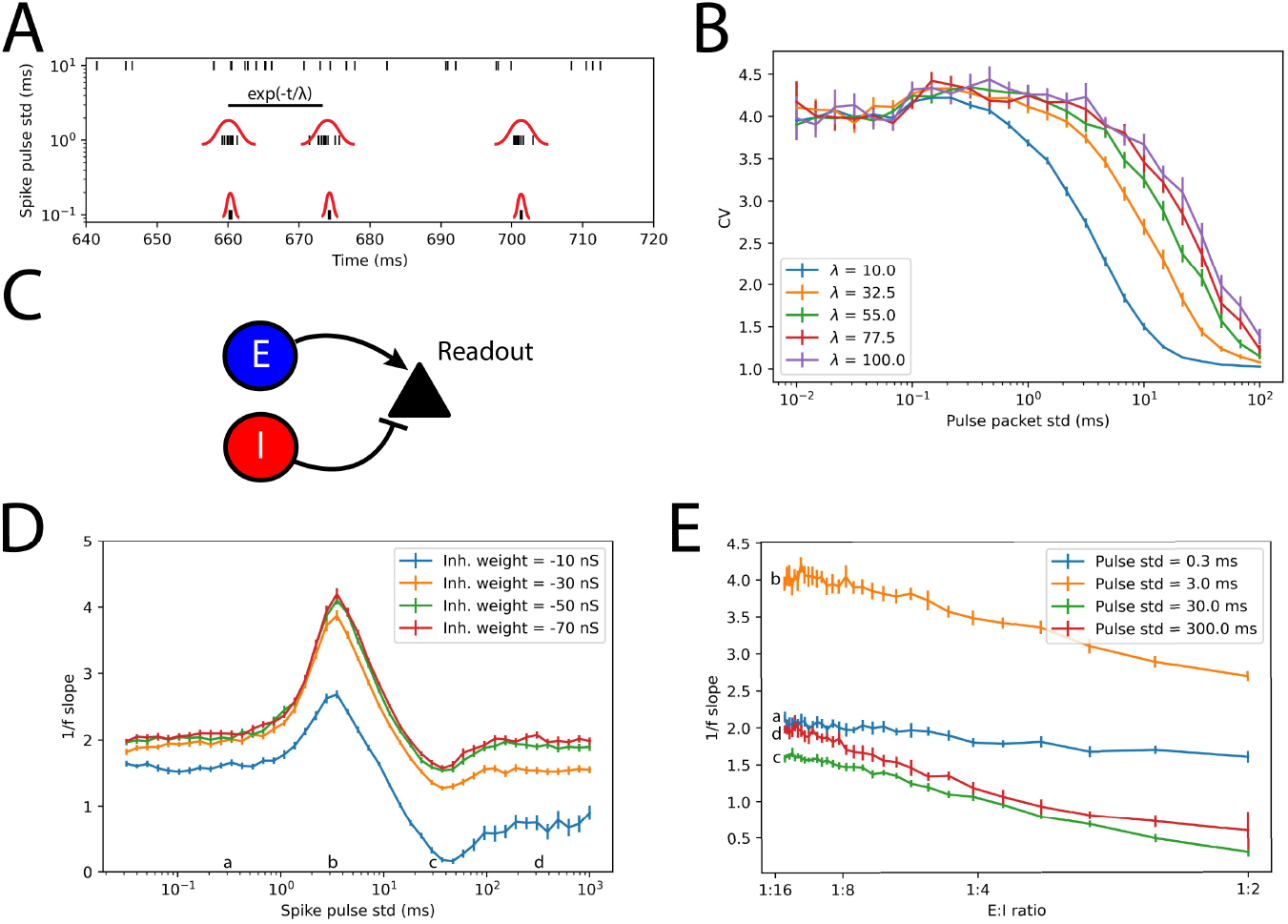
Large values of 1/f slope are only possible at intermediate levels of spike synchronization. (A) A so-called “Gaussian packet model” (GPM) was created by superimposing upon a Poisson process with rate λ a cluster of *N* spikes with a given standard deviation, *σ*. Three examples with identical Poisson processes but successively larger *σ* are shown, with Gaussians drawn schematically in red. (B) A numerical demonstration that for large enough *σ*, the GPM converges to a Poisson point process as indicated by a coefficient of variation (CV) of the inter-spike intervals approaching 1. (C) Spike trains generated by the GPM were assigned to excitatory (E, blue) and inhibitory (I, red) synapses on a Hodgkin-Huxley point neuron (black), called the readout neuron. The 1/f slope of the resulting transmembrane current was measured. (D) The 1/f slope from (C) is shown as a function of *σ*. Different colors denote increasing the inhibitory synaptic weight for a constant excitatory weight of 5 nS. (E) The 1/f slope as a function of E/I synaptic weight ratio. Lowercase letters on the *x*-axis of (D) correspond to traces with the same lower case letter. Each trace corresponds to a 10-fold difference in *σ*. Only for 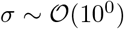 ms leads to 1/f slopes > 2. All error bars indicate 1.96 standard error of the mean.

In words, even though the underlying process is not Poisson, compared against the slower membrane time scale, the highly synchronized pulse of *N* independent spikes coarse-grains to a homogeneous Poisson process of single spikes each with an amplitude scaled by *N*.

On the other end, if *σ*_0_ ≫ λ, the GPM can be thought of as a slowly undulating inhomogeneous Poisson process, or even just a Poisson process with a new rate λ_0_ > λ given by the superposition of N independent point process [Deger et al., 2012], which in the limit of *N* → ∞ can be shown to converge to Poisson statistics. Empirically, even modest *N* still leads to Poissonian properties [Deger et al., 2012]. See Fig. 1B for a numerical demonstration of the convergence of the inter-spike interval coefficient of variation (CV) to 1 as *σ*_0_ is increased, indicating convergence to a Poisson process. In the middle, where *σ*_0_ ≈ *τ_m_*, the GPM resembles some point process that cannot be approximated as a homogeneous Poisson process. The difference between these regimes is illustrated in Fig. 1A.

The GPM spikes were assigned to E and I synapses and fed into a model neuron (Fig. 1C), the resulting 1/f slope of the transmembrane current PSD was measured as a function of the packet width *σ* (Fig. 1D) and the E/I synaptic weight ratio (Fig. 1E). It is apparent that for 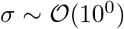 ms, the 1/f slope attains a maximum regardless of the E/I ratio. This range of *σ* corresponds to when the CV is significantly different from 1 (Fig. 1B). The magnitude of the maximum value of the 1/f slope attained in this range of *σ* depends weakly on the E/I ratio — it is clear that at no other values of *σ* can the 1/f slope exceed ~2 just by changing the ratio of synaptic weights (1E). Taken together, these simulations suggest that deviation form a Poisson process, namely by clustering of spikes with a width *σ* ≈ *τ_m_*, leads to extreme values of 1/f slope that cannot be explained by E/I ratio alone.

### Extreme values of 1/f slope in a recurrent network depend on synchronization alone

The GPM is an artificial stimulus and does not represent exactly how background activity in a real biological network would appear. To address how the 1/f slope varies in a more realistic setting, a network of 1,000 (800 E, 200 I) recurrently coupled spiking neurons is simulated. Importantly, the transition between an asynchronous state [Boustani and Destexhe, 2009] (Fig. 2A) and a synchronized oscillatory state (Fig. 2B) is explored.

**Figure 2:**
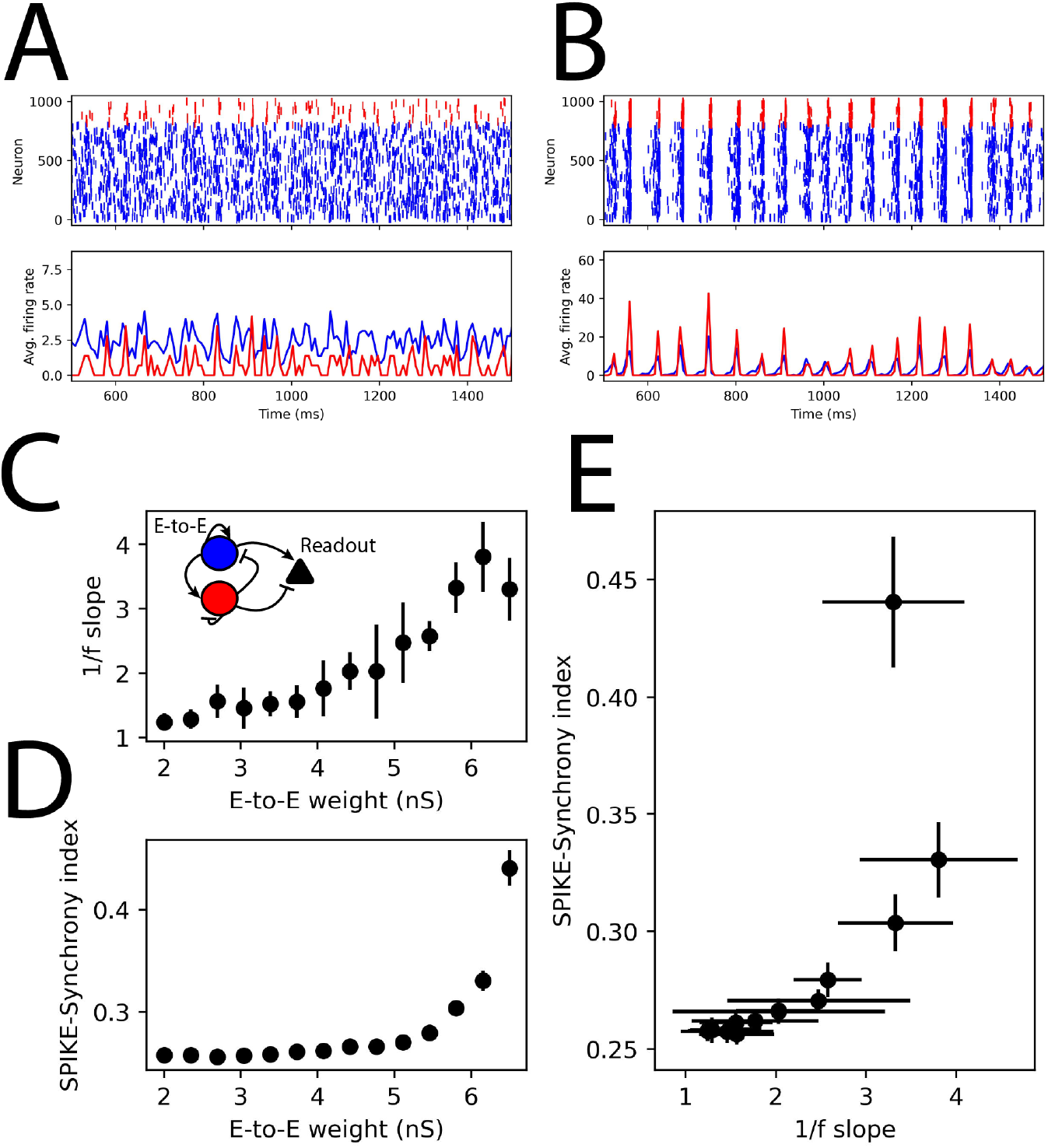
Large values of 1/f slope in a recurrent spiking network correspond to large values of the SPIKE-synchrony index. (A) A spiking recurrent network model in the asynchronous and (B) the synchronous regimes after increasing the E-to-E weight, *w_EE_*, from 3.5 nS to 6.0 nS, respectively. All other network parameters were held constant. (C) The 1/f slope is shown as a function of *w_EE_*. The inset shows *w_EE_* annotated on a schematic of the network. (D) The SPIKE-sychrony index [Kreuz et al., 2013, 2015a,b] as a function of *w_EE_*. (E) The 1/f slope and SPIKE-sychrony index plotted together in the plane. All errorbars indicate 1.96 standard error of the mean.

The control parameter to transition between asynchronous and synchronous activity is the recurrent excitatory weight (E-to-E weight, *w_EE_*). As *w_EE_* increases, so does the 1/f slope and the SPIKE-Synchrony index [Kreuz et al., 2013, 2015a,b] (2C-E). To control for the emergence of a ~ 20 Hz rhythm, peaks in the power spectrum are found and subtracted before estimating the 1/f slope [Donoghue et al., 2020b] (see the Methods for details). Therefore, the correlation between 1/f slope and SPIKE-Synchrony does not depend on the emergence of this oscillation itself, only on the synchronization of spiking activity in this regime. Furthermore, important to note is that this modulation of the 1/f slope does not depend at all on changing the E/I ratio at the readout neuron’s synapses as those weights are left unchanged. Thus this mechanism of generating large 1/f slopes is not explained by the models in Gao et al. [2017] or Trakoshis et al. [2020].

## Discussion

An explanation for steep (> 2) 1/f slopes has been put forth which shows steep 1/f slopes only exist for spike trains which deviate from a Poisson process (CV ≠ 1, see Fig. 1A-B), or from those which cannot be coarse-grained to a Poisson process. Concretely, intermediate synchronization of spike arrival times in a pulse of activity generated by a GPM with standard deviation *σ* ~ 10^0^ ms is sufficient to generate large 1/f slopes in simulated transmembrane currents (Fig. 1C-E). This finding was reproduced when the readout neuron was instead stimulated by asynchronous or synchronous activity generated by a recurrent spiking network model (Fig. 2A-D). Higher degrees of spiking synchronization resulted in larger magnitude 1/f slopes (Fig. 2E), recapitulating the findings with the GPM stimulus of intermediate *σ*.

Prior models concluded that the E/I balance is an important driver of variation of the 1/f slope [Gao et al., 2017, Trakoshis et al., 2020]. Several recent experimental studies also linked biological processes thought to result in alterations of E-I balance like aging [Schaworonkow and Voytek, 2021], ADHD and medication [Robertson et al., 2019], and autism, gender, and E & I gene expression maps [Trakoshis et al., 2020] to argue in-favor of an E-I (im-)balance hypothesis in different brain states. This hypothesis has been successful for 1/f slopes < 2. In contrast, the present study suggests that E/I balance alone does not directly result in the observed 1/f slope slopes >2. Instead, changes in the underlying spike timing statistics of the microcircuit play a larger role when 1/f slopes are very steep. Therefore, when 1/f slopes larger than 2 are present, one must consider the possibility of non-Poisson spiking statistics in the network. This is consistent with recent modeling work by Wardak and Gong [2021] who found that only slight deviation of spiking statistics from unity CV (about 1.5 in the “fractional” regime) still persevered 1/f slope ~ 2, suggesting large deviation from Poisson statistics are required for slopes > 2.

Before discussing potential applications of these findings, first several limitations of the present model must be discussed. Firstly, the model only considers sub-threshold activity—i.e. the readout neuron is assumed not to spike. Second, the E/I ratio is only tested based on prior literature that the ratio by itself explains variability in the 1/f slope [Gao et al., 2017], so it may be that more careful exploration of the full E-I weight plane may uncover regimes where our findings are not valid. Also, the model neuron is considered to be a point source, however, it is known that spatially extended neurons may have different 1/f slope along their extent [Gao et al., 2017]. Finally, the stimuli were intended to model only resting/background activity, so the effects of a strong, time varying driving stimulus are not considered. To build models of complex neural activity using this framework, these limitations of the present study must be examined.

The present study is novel because it extends the analysis of 1/f slopes to highly synchronous brain activity. This model opens the door to use 1/f slopes to infer latent spike synchronization statistics, potentially during burst-suppression from anesthesia and coma [Steriade et al., 1994, Amzica, 2015, Muhlhofer et al., 2017, Kenny et al., 2014, Lewis et al., 2013, Shanker et al., 2021, Ferron et al., 2009, Kroeger et al., 2013]. For example, Kenny et al. [2014] demonstrated propofol and sevoflurane have different burst-suppression patterns on EEG, and thus 1/f slope may be useful in further classifying these bursting patterns. Likewise, 1/f slope might also be used to differentiate or prognosticate coma status in parallel with existing burst suppression metrics, or even perhaps guide pharmacological interventions on coma. For example, the time to resumption of EEG bursting after cardiac arrest depends on cerebral perfusion [Crouzet et al., 2016, Azadian et al., 2021, Wilson et al., 2019], so 1/f slope analysis of this bursting state might constrain microcircuit models of how re-perfusion modulates neuronal dynamics. Thus, the present model of 1/f slope may be used to further understanding of how different anesthetics modulate the synchronization of neural circuits and subsequently alter consciousness.

## Methods

### Gaussian packet model

All simulations were done using NEST version 3.1 [Deepu et al., 2021]. To create the Gaussian packet model (GPM) simulations, the NEST generator `pulsepacket_generator` was used with specified standard deviations (listed in Figure 1) and inter-pulse times Δ*t* drawn from an exponential distribution

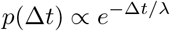

with λ = 50 ms in Figures 1A, D-E, and varied from 10 to 100 ms in Fig. 1B. The excitatory GPM was set to emit 10 spikes per packet, and the inhibitory was set to 20 spikes per packet. The simulation was run for a total of 5 s with 30 replicas at each setting of *σ*. Note, the default time grid spacing of 0.1 ms was kept for all simulations. Modifying this value to 0.01 ms made no difference to the results when using the GPM.

### Model readout neuron

The readout neuron was a NEST `hh_cond_beta_gap_traub` model which simulates a Hodgkin-Huxely neuron [Hodgkin and Huxley, 1952] with bi-exponential synaptic conductances of the form

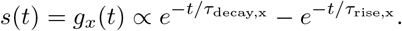

The transmembrane current was defined to be

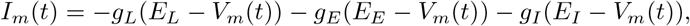

where *V_m_* is the membrane potential, *E_x_* and *g_x_* denote reversal potentials and the leak (*x* = *L*), excitatory synaptic (*x* = *E*), and inhibitory synaptic (*x* = *I*) conductances, respectively. Specific readout neuron synaptic weight settings are given in the corresponding figure legends. All other parameters were kept as NEST defaults except for the membrane capacitance *C_m_* = 150 pF, the inhibitory reversal potential *E*_I_ = −80 mV, *τ*_rise,E_ =0.1 ms, *τ*_decay,E_ = 2 ms, *τ*_rise,I_ = 0.5 ms, *τ*_decay,I_ = 10 ms, the latter synaptic parameters chosen to be consistent with Gao et al. [2017].

### Recurrent spiking network model

800 excitatory and 200 inhibitory NEST `iaf_cond_alpha` neurons were simulated, with default “alpha” synaptic conductance pulses. The membrane potentials were initialized at −62 mV with standard deviation 1.5 mV. All the neurons received a constant external current *I_e_* of mean 100 nA, standard deviation 15 nA if they were E, mean 140 nA, standard deviation 15 nA if they were I. Each of the E neurons were stimulated with a Poisson generator with rate of 140 Hz and synaptic weight of 14 nS. The recurrent synaptic weights are given in Table 1. The neurons were randomly connected with a fixed in-degree of 50 synapses per neuron. The network was simulated for 2500 ms, and the resulting spiking activity of randomly selected 400 E neurons and 100 I neurons were sent to the readout neuron with E weight of 5 nS and I weight of −20 nS (E/I ratio of 1/4).

**Table 1:** A list of recurrent synaptic weights for simulations in Fig. 2

| Connection type | Weight |
| --- | --- |
| E-to-E | 2 to 6.5 nS |
| E-to-I | 10 nS |
| I-to-I | $-3$ nS |
| I-to-E | $-3$ nS |

### Measuring 1/f slope and SPIKE-synchrony

From *I_m_*(*t*) defined above, the FOOOF package [Donoghue et al., 2020b] was used to estimate the 1/f slope. First, the first 500 ms of *I_m_*(*t*) were truncated to remove transients. Then, *I_m_*(*t*) was detrended and then was transformed into a power spectral density (PSD) via Welch’s method (sampling rate = 1000 Hz, Hann window with 400 samples, 300 sample overlap). Then the PSD was passed to FOOOF. For the GPM model, only the aperiodic component was fit between 4 to 100 Hz using the “knee” method. For the network model, a maximum of three peaks were included with `min_peak_height` = 0.5 and `peak_threshold` = 3.

The SPIKE-synchrony index was measured using the pyspike package in Python [Kreuz et al., 2013, 2015b,a]. 10 replicas were used for each measurement of 1/f slope and SPIKE-synchrony index seen in Fig. 2.

Signal processing tasks (detrending, Welch’s method, etc.) were accomplished using SciPy [Virtanen et al., 2020].

## Code availability

Jupyterlab notebooks of the Python code used in all simulations are provided in this [GitHub repository](https://github.com/justidy1/Spike_sync_1f_slope.git): https://github.com/justidy1/Spike_sync_1f_slope

## Acknowledgements

This work was supported by R21EB024793 to Y.A. and the Roneet Carmell Memorial Endowment Fund to Y.A. JDY is supported by a National Institutes of Health Training Grant, T32-GM008602. The content is solely the responsibility of the authors and does not necessarily represent the official views of the NIH.

## Notes

https://github.com/justidy1/Spike_sync_1f_slope

### Competing Interest Statement

The authors have declared no competing interest.

